# Evaluating the Sensitivity of Dry and Gel-Based Wearable EEG for Cognitive Load Estimation

**DOI:** 10.64898/2026.05.05.723048

**Authors:** Sebastian Idesis, Mireia Masias Bruns, Parvin Emami, Saravanakumar Duraisamy, Luis A. Leiva, Ioannis Arapakis

## Abstract

**Purpose:** We present a large-scale (N=120) comparative study of gel-based and dry electroencephalography systems for cognitive load analysis in tasks involving information visualization stimuli. Although dry systems are increasingly adopted owing to their portability and fast setup, their sensitivity to cognitive-related measurements (as compared to gel-based systems) remains debated. This limits the understanding of whether dry systems provide sufficient sensitivity for cognitive load assessment under controlled task conditions.

**Methods:** We analyzed a diverse set of signal quality metrics, such as signal-to-noise ratio and channel retention, combined with spectral features across frequency bands to evaluate the ability for each device to capture workload-related neural markers during information visualization tasks.

**Results:** Although the gel-based device showed consistently better quality results than the dry one, the effect sizes suggest a small practical significance of the differences between systems. These results demonstrate that dry systems can provide adequate physiological sensitivity for cognitive load assessments.

**Conclusion:** Our findings highlight the trade-off between usability (setup, calibration, etc.) and data fidelity, providing practical guidance for choosing electroencephalography systems for cognitive workload monitoring and applied neuroengineering research. Overall, the results suggest that dry systems can support coarse-grained cognitive load assessment, while gel-based systems remain advantageous when greater sensitivity is required.

## 1 Introduction

Electroencephalography (EEG) plays a central role in neuroscience, clinical diagnostics, and neurotechnology applications. Traditionally used in laboratory environments, these recordings usually involve the use of gel-based electrodes, which require meticulous preparation and wired amplifiers, thus ensuring stable, artifact-reduced signal acquisition. The growing interest in portable EEG systems for real-world deployments [1] has led to advancements in dry electrode technologies. While offering a more user-friendly alternative -reducing the preparation time and facilitating broader adoption [2]-this comes at the expense of higher impedance and increased susceptibility to motion artifacts. However, the practical trade-offs between dry and gel-based EEG remain insufficiently characterized. Most existing studies have focused on signal quality, inter-system agreement and usability [3, 4], whereas others have examined their sensitivity and robustness in distinguishing overt pathological conditions from healthy states [5]. However, analyses of subtle variations in cognitive or mental activity remain comparatively rare and, rely on limited sample sizes [6]. High-fidelity recordings are essential for reliably capturing such nuanced neural differences, making systematic comparisons between dry and gel-based systems crucial for understanding their respective strengths and limitations [7], particularly important in applications such as field-based cognitive monitoring, neuroergonomics, and ambulatory neural states [1]. In this study, we address these gaps by evaluating the reliability of both gel-based and dry EEG systems in capturing task-induced cognitive load, operationalized through differences between task difficulties in an Information Visualization (InfoVis) context [1].

In recent years, EEG has become a popular method for assessing cognitive load across a wide range of task domains, including in data-rich and visually demanding environments. For instance, Murtazina and Avdeenko [8] explored EEG-based indicators to monitor cognitive load in intelligent learning systems, demonstrating the potential of real-time neural metrics for adaptive cognitive assessment. In another study, Anderson et al. [9] used a dry-electrode EEG system to assess cognitive load across various statistical visualization designs, providing objective evidence beyond self-report measures.

Similarly, Nuamah et al. [10] employed EEG to examine how task structure influences cognitive effort, illustrating how EEG-derived metrics can sensitively track mental workload. These studies highlight EEG’s capacity to capture nuanced mental states such as attention, engagement, and workload during visualization tasks. Such evidence underscores the broader value of EEG as a tool for quantifying cognitive load in visually demanding contexts, which is directly relevant for benchmarking the performance of different EEG acquisition systems.[11].

Despite the growing interest in using EEG to assess cognitive load in visually demanding tasks, existing studies rarely address the impact of EEG hardware on the quality and interpretability of neural metrics in such settings. Most prior work relies on either gel-based or dry systems in isolation, without systematically comparing their performance across varied visual stimuli and task demands. Furthermore, little attention has been paid to how visualization complexity and design variations might interact with EEG signal quality, potentially limiting the generalizability of findings across setups and contexts. From a biomedical engineering perspective, understanding these trade-offs is essential for determining whether portable dry-electrode systems can provide the signal fidelity required for reliable cognitive-load estimation in real-world or ambulatory environments.

In this context, we formulated the following research questions:

**RQ1:** How do gel-based and dry EEG systems differ in their sensitivity to cognitive load estimation under controlled task conditions?

**RQ2:** What are the implications of these differences in the use of portable EEG systems for cognitive monitoring and neuroengineering applications?

To address these questions, this study compares one dry and one gel-based EEG system under controlled conditions involving visually complex tasks. We recruited a large and diverse sample of participants who engaged with a broad set of visual stimuli spanning multiple levels of complexity and task demands. Our experimental design incorporates both objective behavioural measures (e.g., accuracy, response time) and EEG-based cognitive metrics to evaluate system performance and to quantify how device characteristics influence the detection of workload-related neural signatures.

Specifically, our approach provides empirical evidence by contrasting one dry and one gel-based EEG system under realistic information visualization scenarios with a relatively large participant pool (N=120), going beyond prior single-device studies. Our contribution is primarily methodological: (1) methodological insights into the strengths and weaknesses of dry versus gel EEG for measuring cognitive load across tasks that require sustained visual processing, and (2) practical guidance for researchers in neuroengineering, cognitive monitoring, and applied EEG research. Rather than proposing visualization design guidelines, the study focuses on system-level differences in EEG sensitivity, which are directly relevant for researchers planning cognitive load experiments using electrophysiological measures, and for determining whether portable dry-electrode systems can offer sufficient fidelity for real-world cognitive assessment [7]. The remainder of this paper is organized as follows. Section 2 reviews the related work. Section 3 describes the experimental design, participants, EEG systems, and analysis procedures. Section 4 presents the results. Section 5 discusses the findings and their implications. Finally, Section 6 concludes the paper.

## 2 Related Work

This section was based on a targeted narrative literature review. Relevant papers were identified through searches in major scholarly databases using combinations of keywords related to dry electrodes, gel electrodes, cognitive load, electroencephalography, information visualization, and human-computer interaction. We prioritized peer-reviewed studies that were directly relevant to signal quality, workload sensitivity, functional connectivity, and practical usability. Additionally, we included a limited number of foundational or closely related references to provide methodological context.

Many prior studies have conducted comparative analyses of EEG recordings using consumer-grade dry-electrode devices and medical-grade gel-based systems [12–14]. Most of these investigations were carried out in healthy populations [7, 15–17], whereas only a few have focused on neurological patients [4, 18]. The findings generally suggest that dry-electrode systems can reliably capture low-frequency EEG components, but often exhibit limitations in signal quality, and particularly in higher frequency bands. However, these comparisons have rarely evaluated whether such differences translate into varying sensitivity to cognitive load, especially under task conditions in which neural modulation is subtle. For instance, Ehrhardt et al. [5] reported that dry EEG systems have a lower Signal-to-Noise Ratio (SNR) and require the rejection of more channels and trials due to artifacts. Nevertheless, these systems have proven capable of detecting Event Related Potential (ERP) components and assessing resting-state connectivity. For Human-Computer Interaction (HCI) applications, this implies that dry systems may be sufficient for detecting stationary neural states but their reduced fidelity at higher frequencies could compromise more fine-grained cognitive measures, such as those involved in distinguishing different levels of task-related cognitive load [19].

Recent research has expanded the evaluation of EEG systems to include their performance in cognitive tasks, often using Deep Learning (DL) classification techniques [20]. These studies demonstrate that gel-based systems typically offer higher performance in detecting subtle EEG patterns, such as those involved in the classification of mental states. This reinforces the view that device selection is not only a technical issue but also determines the granularity at which user states can be detected and acted upon in interactive environments.

Other studies have integrated EEG-based neurophysiological measures into the evaluation of data visualizations. Anderson et al. [9] used EEG to quantify cognitive load associated with different types of graphical representations, validating EEG as a scalable and real-time metric for Information Visualization (InfoVis) assessment. Similarly, Nuamah et al. [10] compared graphical versus numerical formats and showed that graphical visualizations reduced subjective workload, and improved both accuracy and response times, when combined with EEG metrics. In contrast to these works, our study does not aim to evaluate visualization effectiveness. Instead, InfoVis stimuli serve as controlled task conditions for eliciting varying levels of cognitive load to compare the sensitivity of gel- and dry-based EEG systems.

Building on this, Yoghourdjian et al. [21] applied EEG to assess cognitive load during complex network visualization tasks. Using a dry EEG system, the authors found that task difficulty correlated with increased theta power. These results support the use of physiological measures to evaluate perceptual scalability in InfoVis settings. Together, these studies establish EEG as a valuable tool for InfoVis evaluation; however, they did not address how differences in hardware might affect the interpretability of the results.

In light of these findings, it is critical to consider the intended application when selecting an EEG system. Prior device comparisons did not examine how these hardware differences affected the detection of cognitive load differences across task conditions. As summarized in Table 1, gel-based systems generally outperform dry systems in terms of artifact susceptibility, signal stability, and user preference, although they often involve longer preparation times and are associated with increased participant discomfort. On the other hand, dry systems offer greater usability and portability, but may suffer from higher noise levels and reduced reliability.

**Table 1:**
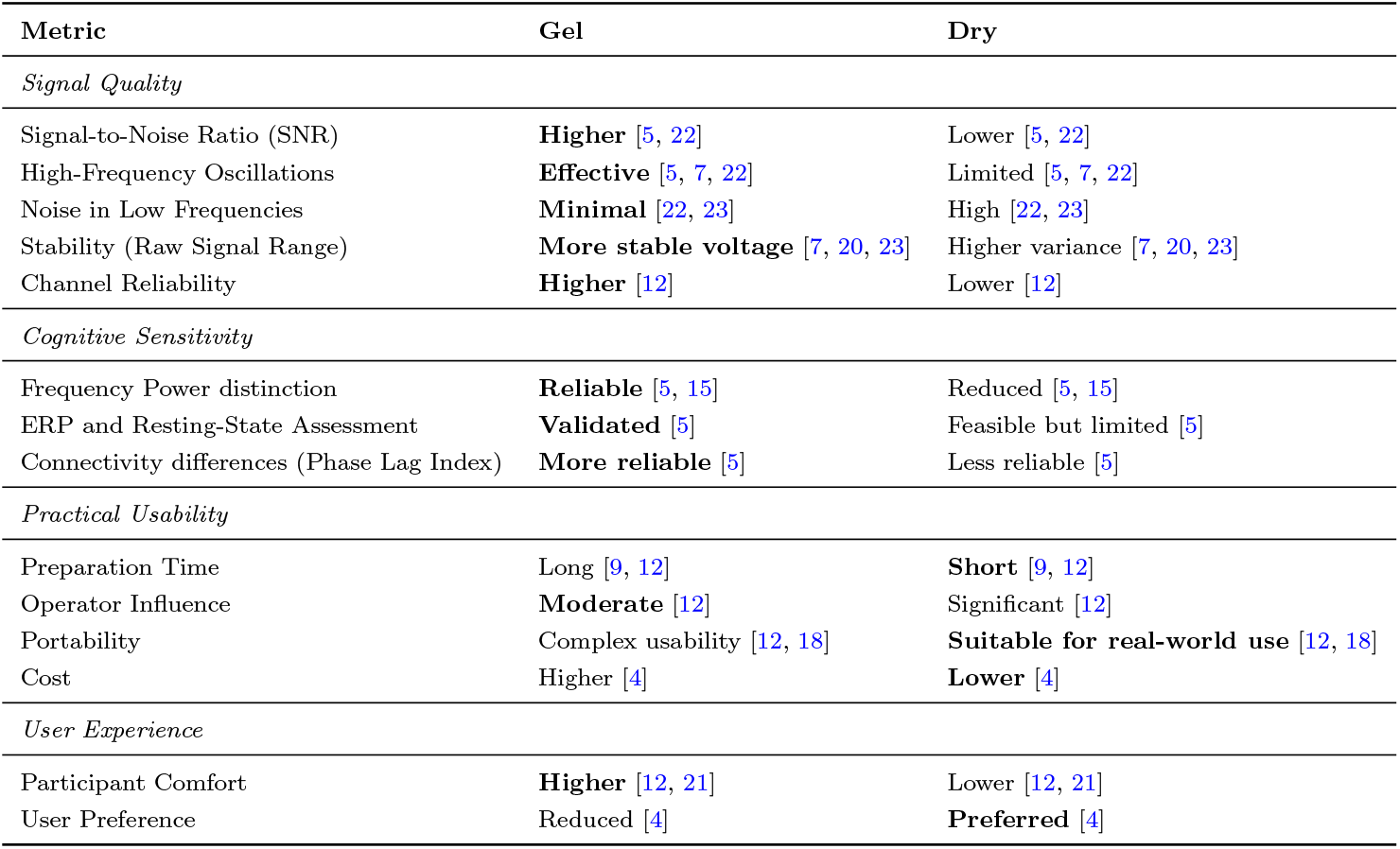
Comparison of Dry vs Gel EEG Systems. Preferred option for each metric is highlighted in bold.

While the comparison of EEG devices has been previously addressed in the literature, these evaluations have not accounted for the impact of different visualization modalities. To our knowledge, few studies have examined EEG signal quality across both sensor types and diverse information visualization formats. By directly linking device-level differences to visualization evaluation, our work advances both the methodological toolkit available to HCI researchers and the practical guidance needed for integrating EEG into user-centred design and adaptive interfaces.

## 3 Methods

We conducted a controlled experiment comparing gel-based and dry EEG systems in interactive visualization tasks, designed to evaluate signal quality and cognitive sensitivity under realistic HCI conditions.

### 3.1 Participants

A total of 120 participants took part in the study. Participants were equally distributed across two research laboratories (each using a different EEG system), with 60 participants recruited per site. Every participant observed a different visualization type, resulting in 20 participants randomly assigned at each site and condition (2 sites × 3 visualizations × 20 participants). Each participant was randomly assigned to only one visualization type to prevent potential learning or exposure effects across conditions.

The participants had normal or corrected-to-normal vision and were proficient in English to ensure consistent task comprehension and response quality. All participants provided written informed consent prior to participation and received monetary compensation for their time. None of the participants reported a history of neurological or psychiatric disorders, nor were they taking any medication that could interfere with the results.

Participants were recruited through mail lists and public advertisements, ensuring a diverse sample beyond the immediate lab environment. The demographics of each participant group are provided in the Tables 2 and 3, including age and gender distribution. Briefly, the overall mean age was 27.89 years (SD = 7.49) and the sample included 49 females and 71 males.

**Table 2:** Age Statistics by Visualization Type and Source.

| **Viz Type** | **Gel-based** |  |  | **Dry-based** |  |  |
| --- | --- | --- | --- | --- | --- | --- |
|  | Graph | Distribution | Timeline | Graph | Distribution | Timeline |
| Mean | 29.5 | 29 | 29 | 26.2 | 26.6 | 27.05 |
| STD | 8.87 | 9.46 | 8.89 | 6.07 | 4.92 | 5.75 |

**Table 3:**
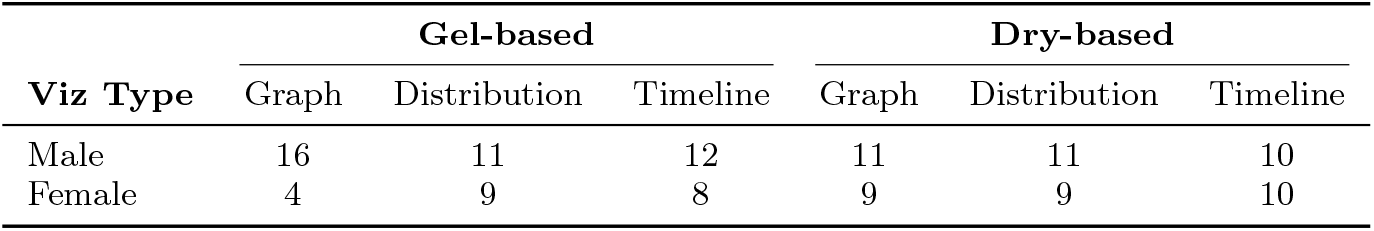
Gender Distribution by Visualization Type and Source.

Finally, the study protocol was reviewed and approved by the ethics committee of the academic participating institution. All procedures complied with the Declaration of Helsinki on ethical standards for research involving human participants.

### 3.2 EEG Systems

#### 3.2.1 Gel-based EEG

For our gel-based Brain-Computer Interface (BCI) device, we utilized a 32-channel actiCHamp system (Brain Products GmbH, Germany). A high-impedance amplifier was used to ensure optimal signal quality and minimal noise. Electrode placement followed the international 10–20 system, the standard in EEG acquisition [24]. The ground and reference electrodes were positioned at Fpz and Cz, respectively. Impedances were kept below 10 kΩ, following standard practices. Electrode preparation took approximately 15 minutes per participant. The list of electrodes is described in Table 4 and shown in Figure 1. A prior study has used the aforementioned device to conduct a quality assessment study [23]. This device has been widely adopted in cognitive neuroscience research and is considered a reference-standard system for high-quality EEG recordings.

**Table 4:** Electrode sets for each system, aligned with Figure 1. Shared electrodes are highlighted in bold.

| Metric | Gel-based | Dry-based |
| --- | --- | --- |
| Channels | **Fp1, Fp2**, F7, Fz, F8, **F3, F4**, FT9, FC5, FC1, FC2, FC6, FT10, T7, C3, C4, T8, CP4, CP1, CP2, CP6, TP9, **P3, P4**, Pz, TP10, P7, P8, **O1, O2**, Oz | **Fp1, Fp2**, AF7, AF8, **F3, F4, P3, P4**, PO7, PO8, **O1, O2** |
| Ground | Fpz | Fpz |
| Reference | Cz | A1 (left mastoid) |

**Fig. 1:**
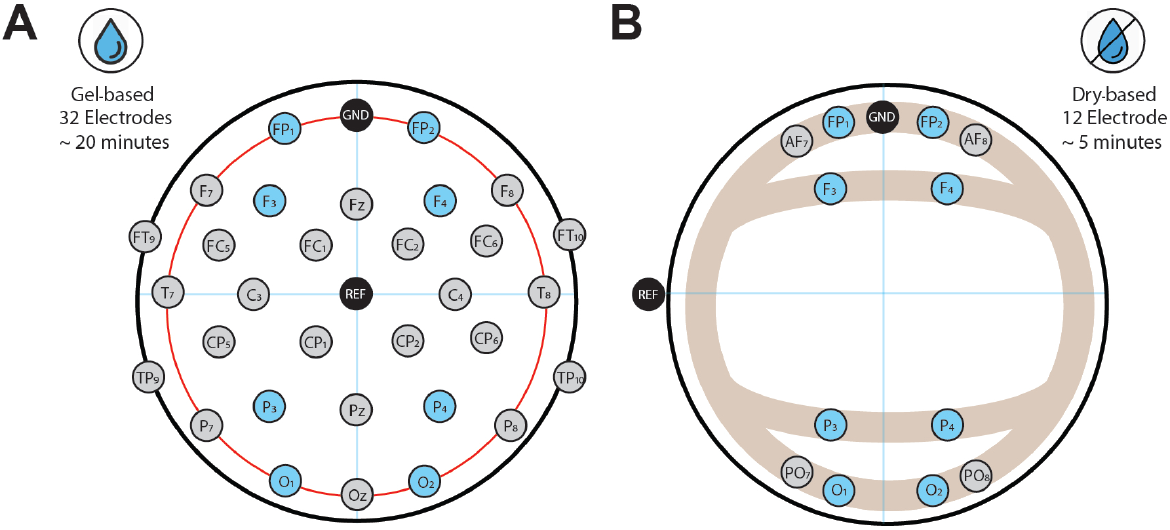
Visualization of the EEG hardware setups used in the study. (A) Gel-based EEG system with 32 channels, including standard 10–20 electrode placement and conductive gel application. (B) Dry EEG system with 12 dry-contact electrodes and reduced setup time. Both setups were used under equivalent environmental conditions across sites. In green, electrodes appearing in both setups. This contrast highlights the trade-off between signal fidelity and usability, which is central to evaluating EEG technology in HCI contexts.

#### 3.2.2 Dry-based EEG

For our dry-based BCI device, EEG data were acquired using a Bitbrain Diadem system, featuring 12 dry electrodes [25]. Electrodes A1 and Fpz were used for reference and ground electrodes, respectively, following the 10–10 system. Raw EEG signals were recorded for subsequent processing. No conductive gel was required. Setup time was approximately 5 minutes, and signal quality was monitored via the system’s internal quality indicators. The list of electrodes is described in Table 4 and shown in Figure 1. Previous studies have used the aforementioned device to conduct a quality assessment study [26]. This system has been specifically designed for applied and field research, prioritizing portability and fast deployment over dense electrode coverage. Because the two EEG systems were recorded at different sites and also differed in channel count, electrode montage, and reference scheme, the present study should be interpreted as a comparison between two complete acquisition setups rather than as an isolated comparison of electrode type alone.

### 3.3 Task design and stimuli

The task was presented on a screen of 1920 x 1080 pixels to ensure consistent resolution across all recordings and participants. The experiment started with the explanation of the task, followed by a set of eight practice trials to assess whether participants correctly understood the task instructions. Once the experimenter confirmed that there were no questions about the task to be performed, the experiment began. The order of the stimuli was randomized across participants to mitigate learning or fatigue effects. Each session consisted of 120 trials, and each trial implemented the following steps (see Figure 2):

**Fig. 2:**
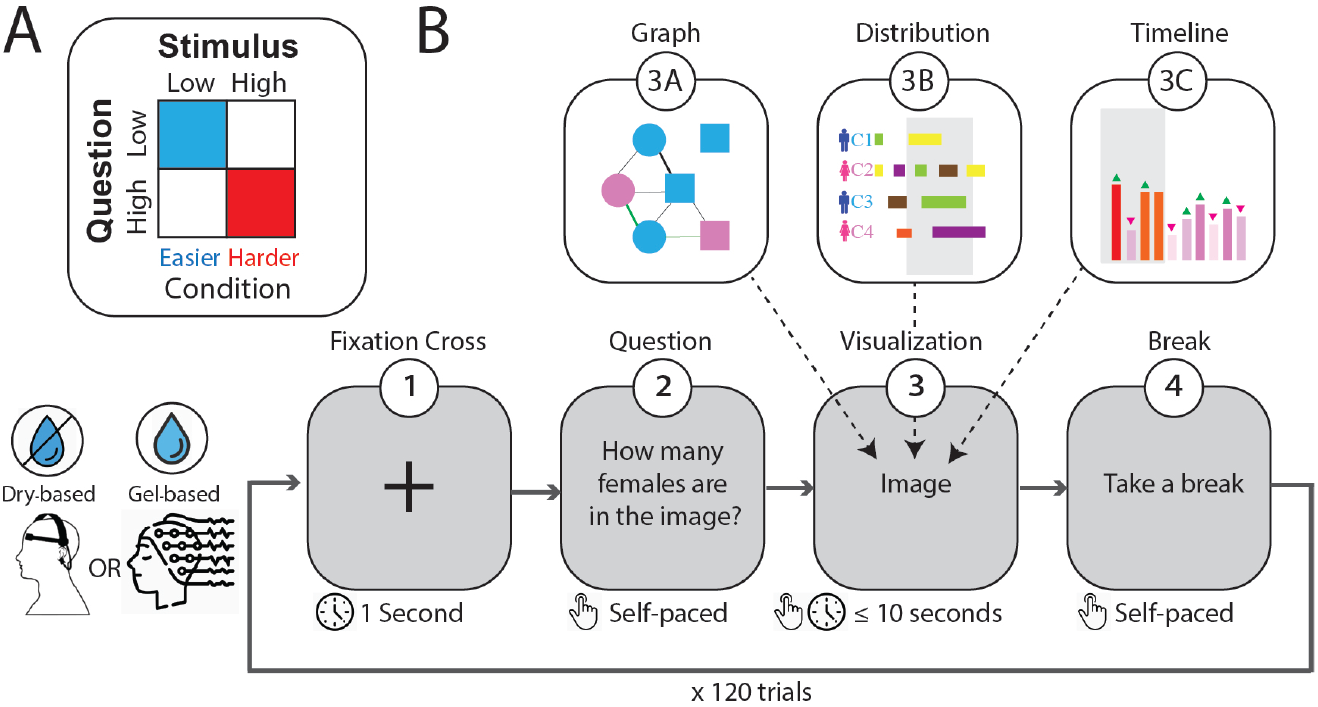
Experimental design. (A) Cognitive load was manipulated through a 2×2 design varying question difficulty (low/high) and stimulus complexity (low/high). In this paper, only the low-low and high-high combinations were compared as Easy and Hard conditions, respectively. (B) Trial structure: each of the 120 trials began with a fixation cross (1s), followed by a self-paced question screen, a visualization (graph, distribution, or timeline; max 10s), and a self-paced break screen. Participants completed the task using either a dry- or gel-based EEG system.

**Fig. 3:**
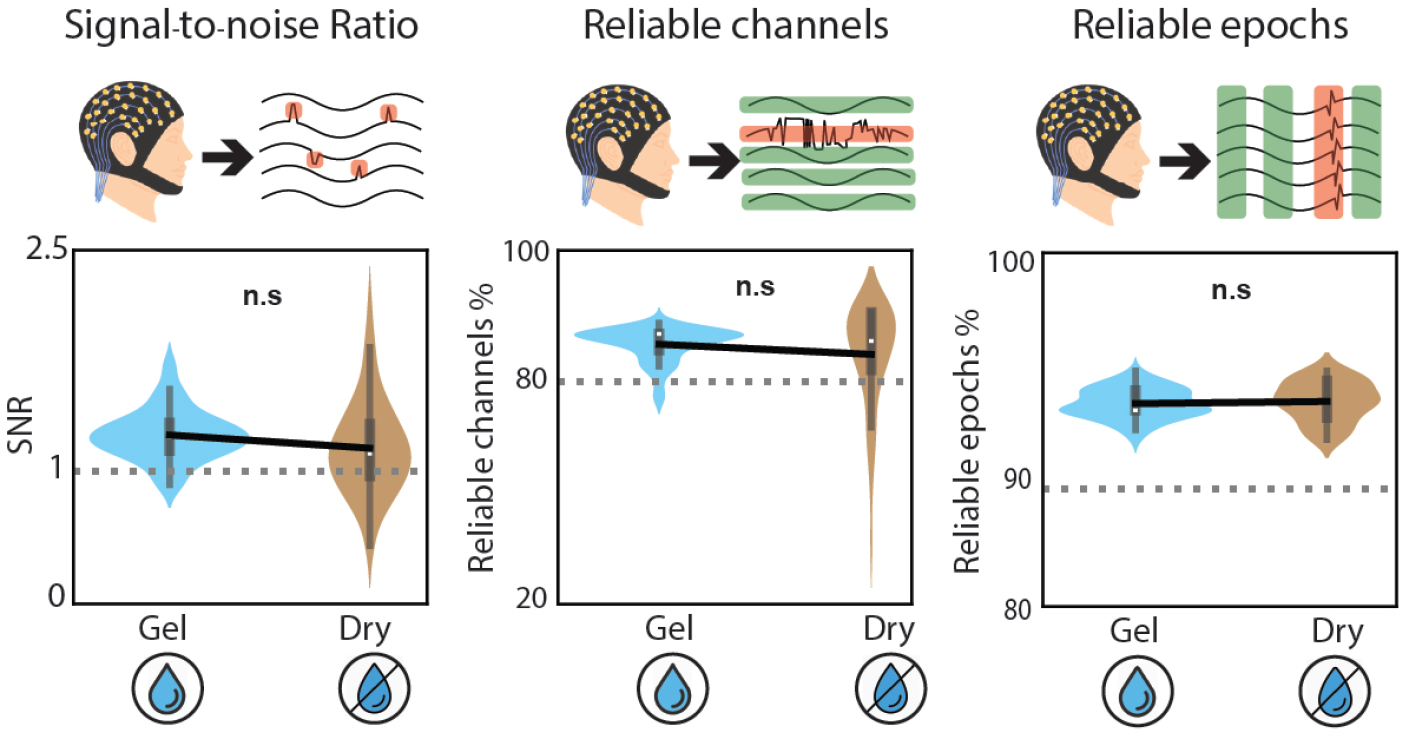
Comparison of signal quality metrics between gel-based and dry EEG systems. Violin plots show signal-to-noise ratio (SNR) (left), ratio of reliable channels (centre), and percentage of usable epochs per participant (right). The gel and dry systems showed comparable signal-quality distributions across the three metrics, with no significant overall differences in SNR, reliable channel ratio, or usable epochs.

1. Fixation cross: Located in the centre of the screen for 1 second.
2. Question screen: Questions about the upcoming visualization. This was self-paced, ensuring that the participants had sufficient time to read and understand the question.
3. Visualization screen: Visualizations including a criminal network information on which the user bases their response. The duration of the screen is 10 seconds or until the subject responds. Responses were restricted to single-digit numbers (0–9).
4. Resting period: Self-paced response screen for the subjects to rest in case it is needed.

The two difficulty factors were defined as follows. Question difficulty was manipulated by adjusting the semantic and numerical reasoning requirements needed to answer each item. Low-difficulty questions involved simple feature detection or direct recall, whereas high-difficulty questions required integrating multiple attributes or inferring relationships within the visualisation. For example, an easy question queried a single attribute (e.g., ‘How many females are shown?’), while a hard question required combining two attributes (e.g., ‘How many females have been arrested more than 10 times?’).

Stimulus complexity was manipulated by varying the amount and density of information in each visualisation. Low-complexity stimuli contained fewer elements, whereas high-complexity stimuli introduced more items and denser structures. Concretely, low-complexity graphs contained three nodes while high-complexity graphs contained six; low-complexity timelines displayed two callers over five hours whereas high-complexity timelines showed four callers over 12 hours; and low-complexity distribution charts covered five hours of call activity while high-complexity versions covered 24 hours.

In our analyses, trials combining low question difficulty with low stimulus complexity were treated as Easy conditions, whereas trials combining high question difficulty with high stimulus complexity were treated as Hard conditions. The mixed conditions (low question difficulty with high stimulus complexity, and high question difficulty with low stimulus complexity) were not included in this binary Easy-versus-Hard comparison. This choice was made to contrast the two most clearly separated task conditions, but it also means that the present analyses do not test the full 2×2 factorial design.

All categories of visualisations in our study show information about a criminal network and the interaction between the members. This design ensured that both behavioral and neural differences could be interpreted with respect to controlled variations in task difficulty. Specifically:

- Graphs: Nodes in the graph represent different individuals with criminal background, while the edges that connect them indicate an association. Graphs also contained information about people’s name, gender, number of arrests, whether they are currently in prison, their birthplace, as well as the strength of the connection with another individual in the network [21, 27].
- Timelines: Timeline visualisations reveal the history of calls between individuals in the criminal network, at different times of the day, indicating the person who is calling (caller), the person that receives the call (callee), and the duration of the call.
- Distribution charts: Distribution chart visualisations indicate the amount (i.e. total number) of calls at a set granularity level; in our case at each hour of the day.

### 3.4 Procedure

The experimental task was implemented using PsychoPy, a platform for stimulus presentation and behavioural response logging [28]. Specifically, PsychoPy presented the visualizations and recorded participants’ answers. To ensure synchronized multimodal data collection across sites, all acquisition devices (EEG and behavioural logging) were time-aligned using a shared timestamp protocol. Said devices operated on the same local network, maintaining consistent internal clocks and allowing for precise temporal alignment across all data streams. This setup ensured that behavioural events and EEG recordings could be matched with millisecond-level accuracy, a requirement for subsequent cognitive workload analysis.

### 3.5 Data acquisition and preprocessing

For both EEG devices, signals were sampled at 256 Hz, providing adequate temporal resolution for downstream analysis. Furthermore, recordings took place in a sound-attenuated, electrically shielded room to minimize external interference. Particular attention was given to artifact-related contamination, including ocular, muscle, motion, and electrode-contact artifacts, through visual inspection, filtering, independent component analysis, and the exclusion or interpolation of noisy channels where appropriate. With equivalent pre-processing, the results obtained from both devices could be compared.

#### 3.5.1 Gel-based EEG

Before recording, the scalp was cleaned and mildly abraded to reduce impedance. Visual inspection ensured all electrodes were properly attached, and impedance levels remained within acceptable thresholds throughout the session. Moreover, artifact removal was carried out using Independent Component Analysis (ICA), filtering any ocular or muscle-related components present in the EEG data. Channels contaminated with persistent noise were interpolated using spherical splines when necessary. These preprocessing steps followed standard best practices in EEG research to maximize data quality and ensure comparability across participants [29].

#### 3.5.2 Dry-based EEG

Electrode contact was verified using the system’s built-in quality indicators, and no skin abrasion or gel was required. Setup time was shorter, typically under five minutes. Preprocessing steps mirrored those applied to the gel-based data. Channels flagged as noisy or unstable were excluded from analysis or interpolated where appropriate. This parallel preprocessing ensured that differences in results reflected device performance rather than differences in signal processing [29].

### 3.6 Data analysis

Offline preprocessing was performed in Python [29], including 50 Hz notch filtering, band-pass filtering between 0.5–40 Hz (FIR filter), artifact rejection, and epoching. Baseline correction was applied using the 500 ms pre-stimulus interval. Because the task was self-paced, epoch durations were not fixed but instead adapted to each trial’s response window. As a consequence, the resulting epochs differed in duration across trials and conditions, which should be considered when interpreting power spectral density, band-ratio, and connectivity measures, since these may also be influenced by differences in temporal structure.

Spectral characteristics were analysed through Welch’s method [30], and log-transformed Power Spectral Density (PSD) values were extracted across canonical frequency bands: delta (1–4 Hz), theta (4–8 Hz), alpha (8–13 Hz), beta (13-30 Hz), and gamma (30–40 Hz) [31]. These features served as the primary markers for assessing cognitive load differences between conditions and comparing system sensitivity.

#### 3.6.1 EEG Feature extraction

In line with methodologies employed by Kleeva et al. [7], our study assessed signal quality using metrics such as SNR across different EEG systems and conditions. Specifically, SNR was calculated through the given formula [5]:

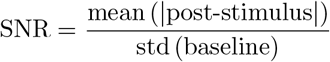

while the reliability of channels and epochs was calculated through MNE-Python pre-defined functions [29].

This SNR measure was used here as a relative descriptive index of signal quality following prior EEG-comparison work, rather than as a standardized quality metric specifically validated for self-paced visual reasoning tasks [5, 7].

In addition, following Ehrhardt et al. [5], our study utilised both time-domain and frequency-domain analyses to evaluate the performance of dry and gel-based EEG systems in detecting cognitive features and assessing functional connectivity (see Appendix for a formula of the metric). For example, as indicated by Yoghourdjian et al. [21], we used EEG-derived power at frontal electrodes to evaluate cognitive workload during visual analytics tasks. Their work highlights the suitability of dry EEG caps for such paradigms, while acknowledging noise-related limitations. Another study by Anderson et al. [9] analysed cognitive workload using band activity at pre-frontal sites. Their use of a low-cost dry EEG headset to measure extraneous load supports the viability of lightweight systems for real-time cognitive load estimation in structured visual tasks. Building on these prior approaches, our study implements a rigorous evaluation of gel and dry EEG systems on both signal quality and sensitivity to cognitive load, providing direct evidence of how device choice influences the reliability of workload-related markers.

## 4 Results

This section reports the comparative results obtained from the gel-based and dry EEG systems across multiple evaluation metrics. We first introduce the behavioral results, followed by the signal quality indicators, including signal-to-noise ratio, channel reliability, and data retention. Next, we assess each system’s ability to detect cognitive load differences using spectral power analyses across canonical frequency bands.

### 4.1 Behavioural results

Behavioural performance showed clear and consistent differences between the Easy and Hard conditions across all visualization types and both acquisition sites. As summarised in Table 5, participants were substantially more accurate and responded markedly faster in the Easy trials. This pattern was highly consistent, with all Easy-condition trials yielding higher accuracy and shorter reaction times than their Hard-condition counterparts. These robust behavioural separations confirm that the experimental manipulation of question difficulty and stimulus complexity produced two clearly distinguishable task regimes. However, because the experiment was self-paced and reaction times differed substantially between Easy and Hard trials, the physiological differences should be interpreted with caution, as part of the observed effects may also reflect differences in trial duration and temporal structure.

**Table 5:**
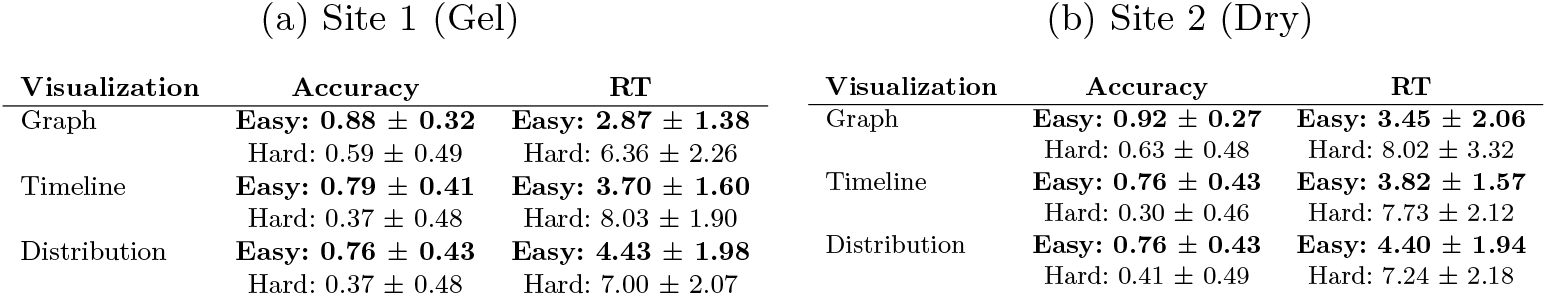
Behaviour metrics for both sites (Gel-based and Dry EEG).

### 4.2 EEG Signal Quality Comparison

To evaluate signal quality, we compared three core metrics across EEG systems, namely (1) SNR, (2) the ratio of reliable channels, and (3) the number of usable epochs per participant. Reference thresholds were included only as descriptive comparison points in the visualisation of signal-quality metrics: SNR *≥*1, reliable channel ratio *≥*80%, and clean epochs *≥*90% of per recording session. Across participants, there was a near-significant difference (*p* =.07) between gel-based EEG SNR values (*M* = 1.23, *SD*= 0.16) and dry recordings (*M* = 1.15, *SD*=.30). The percentage of good channels was also not significantly different (*p* =.17) between the gel-based system (*M* = 90.74, *SD*= 4.98) and the dry system (*M* = 87.81, *SD*= 14.67). Finally, the percentage of artifact-free epochs was also not significantly different (*p* =.54) in the gel condition (*M* = 97.23, *SD*= 1.27) and the dry condition (*M* = 97.41, *SD*= 1.70), indicating similar signal quality.

To evaluate whether differences in EEG signal quality metrics were consistent across visualization formats, we conducted a two-way ANOVA with EEG system (gel vs. dry) and visualization type (Graph, Timeline, Distribution) as factors. For the SNR, we observed a significant main effect of visualization type (*p <*.01), a near-significant main effect of EEG system (*p* =.08), and an interaction effect (*p <*.05). Post-hoc comparisons (Tukey’s HSD) revealed that no significant difference between the conditions (*p >*.11). For the Channel Reliability, neither the main effect of visualization type (*p* =.54), EEG system (*p* =.19), nor the interaction term (*p* =.71) were significant. Last, for the Epoch Acceptance Rate, no significant effects were found: EEG-system (*p* =.56), visualization type (*p* =.35), and their interaction (*p* =.75) was not statistically significant. Altogether, these findings suggest that while the type of visualization influences SNR, particularly with Timeline plots yielding lower values (See Table 6), overall system differences (gel vs. dry) did not significantly impact signal quality metrics, nor did interactions with visualization format. Together, these results indicate that both devices provided recordings of sufficient quality to support subsequent comparisons of cognitive-load sensitivity, while not demonstrating a significant overall difference in the signal-quality metrics examined here.

**Table 6:** Comparison of EEG quality metrics between Gel and Dry datasets for each visualization type and overall. Gel/Dry: mean values; *p*: p-value; *d*: Cohen’s *d*. Bold values indicate Cohen’s *d <* 0.4 (low effect size).

|  | Graph |  |  |  | Timeline |  |  |  | Distribution |  |  |  | Overall |  |  |  |
| --- | --- | --- | --- | --- | --- | --- | --- | --- | --- | --- | --- | --- | --- | --- | --- | --- |
|  | Gel | Dry | *p* | *d* | Gel | Dry | *p* | *d* | Gel | Dry | *p* | *d* | Gel | Dry | *p* | *d* |
| SNR | 1.21 | 1.24 | .75 | **-0.11** | 1.22 | 0.98 | .00 | 1.27 | 1.26 | 1.25 | .87 | **0.06** | 1.23 | 1.15 | .07 | **0.35** |
| Epochs (%) | 97.19 | 97.66 | .38 | **-0.30** | 97.45 | 97.58 | .80 | **-0.08** | 97.06 | 96.99 | .88 | **0.05** | 97.23 | 97.41 | .54 | **-0.12** |
| Channels (%) | 91.61 | 90.10 | .62 | **0.18** | 90.62 | 85.42 | .16 | 0.47 | 89.97 | 88.43 | .73 | **0.12** | 90.74 | 87.81 | .17 | **0.26** |

### 4.3 Frequency Band Power Differences Across Cognitive Load Conditions

We examined power differences across canonical EEG frequency bands (delta, theta, alpha, beta, and gamma) between high and low cognitive load conditions. These conditions were defined based on the amount of visual stimuli presented during each trial. The goal was to assess whether each EEG system could reliably capture neural differences associated with task difficulty, and whether the gel-based system provided greater sensitivity to such effects.

A series of linear mixed-models were conducted separately for each frequency band, with *load condition* (high vs. low) as a within-subject factor and *EEG system* (gel vs. dry) as a between-subject factor. Across the analysed frequency bands, significant main effects of cognitive load were observed in the delta and theta ranges.

A linear mixed-model ANOVA for the delta band revealed a significant main effect of load condition (*p <*.01), indicating that delta power differed between high and low workload trials. There was no significant main effect of EEG system (*p* = 1.00) and no significant interaction between load condition and EEG system (*p* =.37). Post-hoc Tukey’s HSD tests showed that the difference between high and low workload was not statistically significant (Dry: *p* =.15; Gel: *p* =.07) within either EEG system (Figure 4A).

**Fig. 4:**
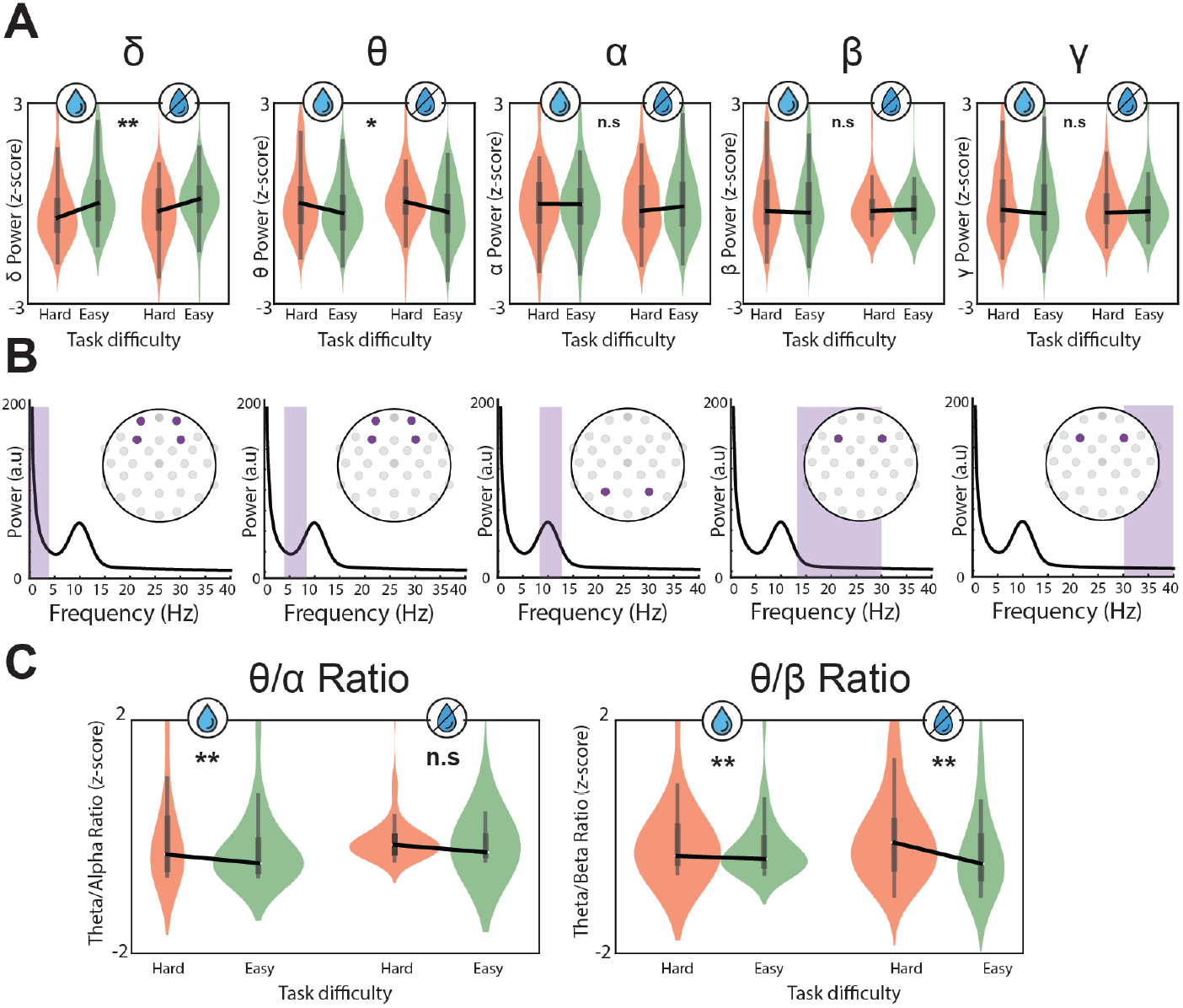
Group-level comparisons of PSD across EEG frequency bands under high and low cognitive load conditions. (A) Violin plots show mean z-scored power in delta, theta, alpha, beta, and gamma bands. Both EEG-devices exhibited stronger task-related modulations, particularly in delta and theta. (B) Electrodes selected for the analysis of each frequency. (C) Comparison between theta/alpha ratio and theta/beta ratio between conditions for both EEG devices.

A similar effect was observed with respect to the theta frequency band showing a main of condition (*p* =.01) but no system or interaction effect (*p >*.5 for both). Post-hoc test showed no difference between the conditions (*p >*.20). For the remaining frequency bands (alpha, beta and gamma), no significant difference were observed in any of the comparisons (*p >* 0.2). However, we note that a more pronounced differentiation was observed when comparing relative proportions across different frequency bands. Specifically, for the theta/alpha ratio, a significant effect (*p <*.01) of workload was observed for the gel-based system, with higher values in the hard condition (*M* = 11.41) compared to the easy condition (*M* = 8.11).

No significant difference (*p* =.5) was found for the dry-based system (hard: *M* = 22.99, easy: *M* = 27.68) (Figure 4C). Values in the axis of the figure were z-scored to facilitate the comparison between the devices. For the theta/beta ratio, both systems showed significantly higher values in the hard condition (Gel: *M* = 16.33, easy: *M* = 10.07; *p <*.01; Dry: hard: *M* = 53.16, easy: *M* = 44.04; *p <*.01) (Figure 4-bottom). These spectral modulations mirrored the behavioural effects reported earlier. However, because the analysed epochs were adapted to each trial’s response window, these differences may reflect not only cognitive-load modulation but also differences in epoch duration and in the relative contribution of perceptual, reasoning, and response-preparation processes across conditions. Overall, these results indicate modest workload-related spectral modulation rather than broad or highly stable differentiation across frequency bands and systems.

### 4.4 Connectivity Patterns Across Cognitive Load Conditions

To explore individual variability in functional connectivity, we calculated the Phase Locking Value (PLV) (For a formula of the metric, see Supplementary Material) between the four frontal electrodes that coincide in both devices (*Fp1, Fp2, F3*, and *F4*). In addition, we computed the mean PLV values for participants in both conditions and compared them in the frequency bands of higher relevance for purely frontal channels (Delta and Theta) (Figure 5A). In both frequency bands there was a consistent significant increase of the PLV for the easy condition (*p <*.01) (Figure 5B). This effect is consistent with the view that lower cognitive demands promote more stable phase alignment among frontal theta generators, as neural resources are not being redistributed toward task-driven updating. Under high load, increased processing demands tend to reduce large-scale synchrony, leading to lower PLV as networks shift toward more flexible and desynchronized patterns to support rapid information integration.

**Fig. 5:**
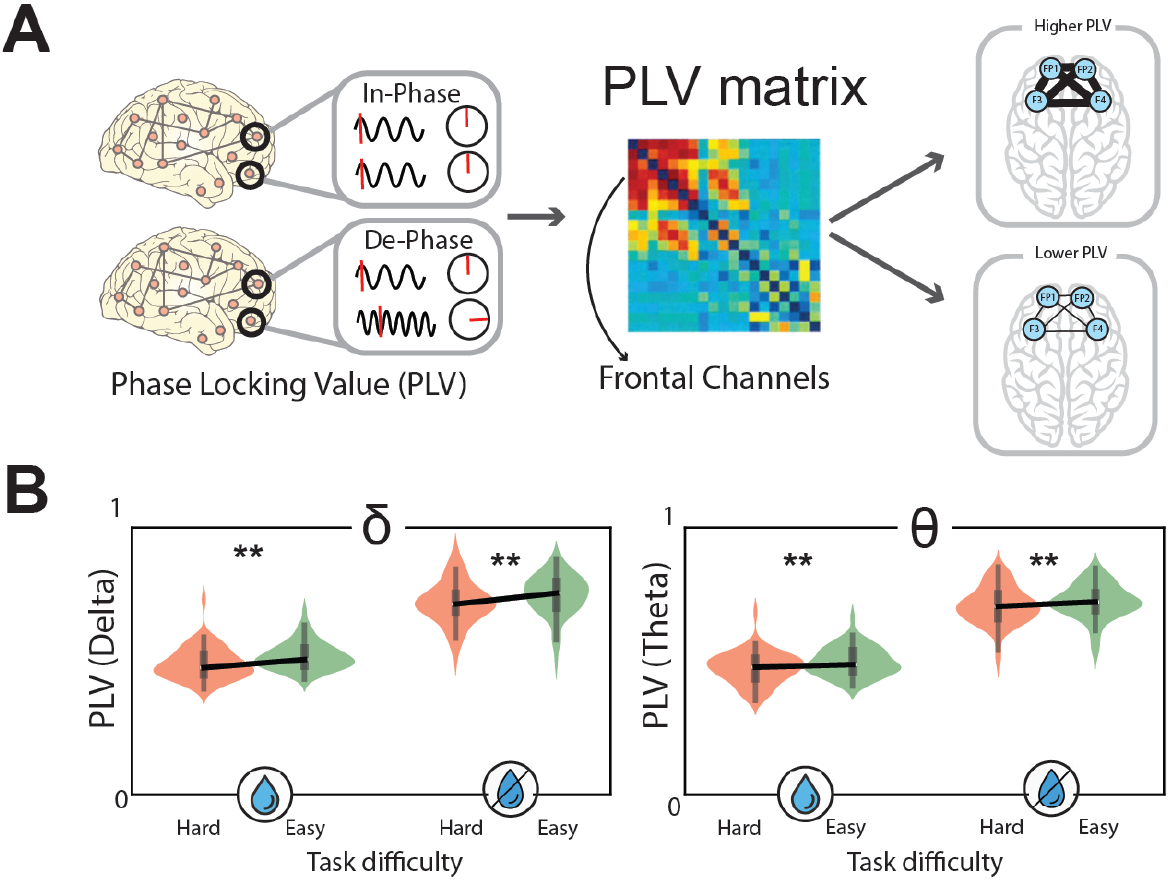
(A) Phase Locking Value calculation through phase difference (in the angles after a Hilbert transformation) between each pair of channels. (B) Comparison of PLV in the different frequency bands of interest (Delta and Theta) for the frontal electrode (Fp1, Fp2, F3 and F4). In all cases, the easy condition elicited a significantly higher level of PLV.

In both frequency bands, the easy condition showed significantly different PLV values than the hard condition. Within the limits of the shared frontal subset analyzed here, this pattern suggests that both systems captured condition-related differences in frontal synchrony. However, because the analysis was restricted to four frontal electrodes shared by both devices and to the delta and theta bands, these results should not be interpreted as evidence of distributed network-level sensitivity. Rather, they indicate that this limited frontal subset was sufficient to detect a condition difference in the present task. Nevertheless, these findings suggest how even revealing significant differences across diverse frequency bands, both devices proved their usefulness to disentangle distinctive mental states. Finally, this interpretation should be considered in light of the self-paced design, since unequal trial durations and differing temporal composition across conditions may also have influenced the connectivity estimates.

## 5 Discussion

This study aimed to evaluate the trade-offs between gel-based and dry EEG systems in the context of cognitive interface research using InfoVis tasks [10]. While both systems achieved comparable signal quality [13, 15, 18, 32], our results demonstrate that gel-based EEG offers only an additional modest sensitivity to cognitive load— particularly under challenging visualization conditions. In line with our study aims, the visualizations served solely as controlled task environments rather than objects of evaluation, and the observed differences reflect device sensitivity rather than visualization design. The following discussion unpacks these findings across key domains: signal quality [13, 15, 18, 32], cognitive sensitivity, practical implications, participant robustness, and future directions.

### 5.1 Signal Quality across systems

Despite the known advantages and superior accuracy of gel-based EEG in controlled lab settings [5, 7, 33], our analysis did not reveal any significant differences in signal-to-noise ratio or channel reliability of epoch retention, suggesting that the performance between gel and dry EEG systems is comparable for our study. This finding suggests that, in the present study, the dry EEG system provided signal quality comparable to the gel system on the metrics examined, although these results should not be interpreted as evidence of equivalence. Therefore, while dry systems generally exhibit higher noise levels and greater variability, their performance remained sufficient for detecting coarse spectral differences between task conditions [7, 8]. This aligns with recent studies showing that dry EEG devices are viable alternatives for certain applications [16], particularly when convenience and speed are prioritized [12, 16]. Importantly, visualization type influenced quality metrics: the timeline plots consistently yielded lower SNR, specially in the dry system, likely due to their higher visual complexity and temporal encoding demands [21]. This convergence in signal quality ensured that subsequent differences in load sensitivity arose from the devices themselves rather than preprocessing or recording artifacts. Importantly, the absence of significant differences in these comparisons should not be interpreted as formal evidence of equivalence between the two systems.

### 5.2 Cognitive Load Sensitivity

Although the signal quality was comparable, the sensitivity of each system to cognitive load variations diverged. Gel-based EEG systems demonstrated somewhat stronger modulation in frequency-band ratios between high and low task difficulty. However, the spectral results were not uniformly strong across bands or systems. In the power spectral density analysis, delta and theta showed significant main effects of load condition, but within-system post-hoc comparisons were not significant for either device. For alpha, beta, and gamma, no significant differences were observed. In addition, the theta/alpha ratio differed significantly only in the gel-based system, whereas the theta/beta ratio showed significant differences in both systems. Taken together, these findings suggest coarse workload-related spectral modulation rather than broad or highly stable spectral differentiation across systems.

Part of the reduced sensitivity of the dry system may be attributable to the lower spatial sampling of the dry system (12 channels) compared to the 32-channel gel-based setup. A denser montage provides broader cortical coverage and more redundancy, increasing the likelihood of capturing workload-related modulations across frontal and parietal regions. By contrast, the sparse dry montage limits spatial information and may attenuate subtle spectral or connectivity effects that rely on distributed sources. Therefore, some of the observed sensitivity differences likely reflect channel-count constraints in addition to differences inherent to electrode type. The same results were observed when inspecting their differences in the connectivity analysis.

The neural modulations aligned with the strong behavioral separation between Easy and Hard trials, supporting the interpretation that both EEG systems captured meaningful cognitive-load variations. Furthermore, the presence of consistent results in the connectivity analysis (Higher PLV for the easier condition) showed the coherent patterns observed in the dry-based and gel-based devices. These neural effects should be interpreted alongside the behavioral results, which consistently showed higher accuracy and faster responses in the easier condition. The alignment between performance differences and EEG-based workload markers indicates that the neural measures reflect meaningful variations in task demands, rather than only capturing differences at the signal level. This coupling between behavioral performance and physiological modulation strengthens the interpretation that the EEG effects reported here are functionally relevant indicators of user experience during visualisation tasks. The fact that both systems revealed the same load-dependent synchrony pattern in this restricted frontal subset supports the view that both devices captured a condition-related difference in the present task, although the narrow electrode selection limits broader network-level interpretation. In conclusion, system performance (beyond raw signal quality) should be the guiding criterion when selecting EEG technology for cognitive interface research.

### 5.3 Implications for EEG-based Cognitive Load Assessment

The present findings have several implications for studies that aim to quantify cognitive load using EEG. First, the consistent behavioural separation between Easy and Hard conditions across all visualisation types provides a clear reference for interpreting the neural effects observed in both EEG systems. The increases in delta and theta power, the modulation of band-ratio measures, and the differences in frontal connectivity align with the performance patterns reported earlier in the Results, indicating that the EEG markers captured meaningful changes in task demand rather than device-specific noise characteristics. Second, the observed load-related patterns suggest that both technologies can detect coarse-grained cognitive load differences in the present task. However, these effects were not uniformly strong across all spectral measures, and the gel system generally showed clearer or more stable modulation in some comparisons. This indicates that dry systems can be used reliably in paradigms where the primary goal is to distinguish broad workload conditions, especially when portability or rapid setup is necessary.

Third, the attenuated sensitivity observed in the dry system highlights an important boundary condition for cognitive load research. Tasks requiring fine-grained discrimination of spectral effects, or analyses relying on distributed cortical patterns, may still benefit from the denser spatial sampling of gel-based devices. The findings therefore emphasise that device choice should be guided by the level of precision required in the cognitive interpretation rather than by signal quality metrics alone.

Overall, these results demonstrate that EEG-based workload assessment is feasible with both systems, but that the depth and granularity of cognitive insights depend on the density and stability of the electrode montage. By clarifying these constraints, the study provides practical guidance for researchers designing cognitive load experiments and interpreting EEG-derived workload indicators.

### 5.4 Limitations and Future Work

An important limitation of the present study is that EEG system type was fully confounded with acquisition site; i.e., each site used their own EEG device. In addition, the two EEG devices differed in channel count, electrode montage, and reference scheme. As a result, the comparison should be interpreted as one between two complete acquisition setups (between-subjects design), rather than as an isolated test of dry versus gel electrodes alone. Accordingly, any differences observed between systems should be interpreted cautiously and should not be attributed solely to the intrinsic sensitivity of dry versus gel EEG technology. Another limitation is that the task was self-paced, resulting in unequal epoch durations across conditions. Because Easy and Hard trials also differed substantially in reaction time, some of the spectral and connectivity effects may reflect differences in temporal structure, and not only differences in cognitive load.

In addition, although the present study focused on highly controlled task-complexity conditions to isolate the neural effects associated with cognitive load, future work should incorporate complementary human-centered measures. Subjective workload ratings (e.g., NASA-TLX), perceived comfort, and qualitative participant feedback would provide additional context to the EEG-based metrics and strengthen the interpretation of user experience. Integrating these measures alongside neurophysiological data would enable a more comprehensive assessment of cognitive demands and enhance the ecological validity of future evaluations.

Furthermore, this study was limited to one gel-based and one dry-based EEG system, which may affect generalizability to other commercial or research-grade devices. Although the task was originally designed as a 2×2 manipulation of question difficulty and stimulus complexity, the analyses reported here focused only on the low-low and high-high combinations. Therefore, the present results should be interpreted as a comparison between two extreme task conditions rather than as a full factorial analysis of the design. In addition, the connectivity analysis was restricted to four frontal electrodes shared by both systems and to the delta and theta bands, which limits the extent to which the PLV findings can be generalized to distributed network-level effects. Regarding the study design, because each participant was assigned to only one visualization type, it reduced potential learning and carryover effects, but it also limits the extent to which visualization-related differences can be attributed solely to stimulus format. Additionally, our analyses centred on spectral features; future work should incorporate ERP-based measures, long-term stability metrics, and possibly real-time classification accuracy. The development of advanced noise reduction pipelines and machine learning-based correction techniques may also help bridge the performance gap between dry and gel systems [20]. Finally, future directions could include testing hybrid systems, integrating eye tracking or physiological markers (e.g., heart rate, pupil dilation), or evaluating cross-device interoperability in adaptive interfaces.

## 6 Conclusion

This study presents a large-scale comparison of dry and gel-based EEG systems in interactive cognitive tasks. Both systems achieved acceptable signal quality and captured some neural differences between Easy and Hard conditions, demonstrating their utility for cognitive-load assessment. While the gel EEG system showed stronger and more stable modulation in some features, dry EEG also detected task-related differences, suggesting that, for the systems and conditions examined here, it may be suitable for studies targeting coarse-grained cognitive effects. Rather than favouring one technology over the other, our results suggest that device choice should be guided by analysis goals and practical constraints such as setup time and user comfort, while taking into account the limitations of the present comparison.

## Statements and Declarations

## Acknowledgments

Research supported by the European Innovation Council through the Horizon Europe’s Pathfinder program (SYMBIOTIK project, grant 101071147).

## Use of Generative Artificial Intelligence

ChatGPT (free version) was used only for language editing and improvement of readability during manuscript preparation. All content was reviewed and verified by the authors, who take full responsibility for the final manuscript.

## Author Contributions

S.I.: Formal analysis, Data Curation, Investigation, Visualization, Writing - Original Draft. M.M.B.: Formal analysis, Data Curation, Investigation. P.E.: Formal analysis, Data Curation, Investigation. S.D.: Formal analysis, Data Curation, Investigation. L.A.L.: Writing - Review & Editing, Supervision, Funding acquisition. I.A.: Writing - Review & Editing, Supervision, Funding acquisition.

## Ethics approval

The study protocol was reviewed and approved by the ethics review board of the University of Luxembourg (study ID: ERP 22-071). All procedures involving human participants were conducted in accordance with relevant guidelines and regulations.

## Consent to participate

Informed consent was obtained from all individual participants included in the study.

