## Supplementary Material for "Evaluating the Sensitivity of Dry and Gel-Based Wearable EEG for Cognitive Load Estimation"

### Appendix

#### Phase-locking value calculation

**Phase-locking value (PLV, theta).** After bandpass filtering $x_c(t)$ to $\theta$ and taking analytic phases $\phi_c(t)$, the pairwise PLV between channels $i, j$ is

$$\mathrm{PLV}_{ij} = \left| \frac{1}{T} \sum_t e^{\,\mathrm{i}\,(\phi_i(t) - \phi_j(t))} \right|.$$

We report the mean across channel pairs:

$$\texttt{PLV mean weight (theta)} = \frac{2}{C(C-1)} \sum_{i<j} \mathrm{PLV}_{ij}.$$

#### Stimuli examples

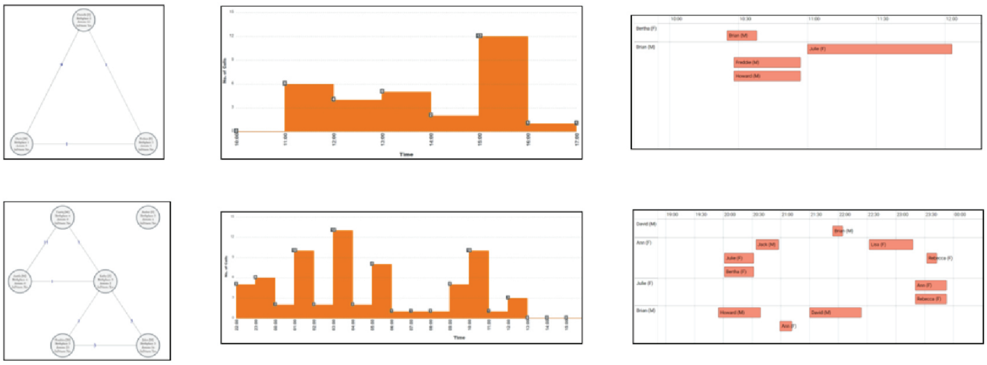

**Fig. 1** Visual conditions visualizations showing low amount of stimuli (top) and high amount of stimuli (bottom) for graph (left), distribution (center) and timeline (right)
